# Impact of pterygium on the ocular surface and meibomian glands

**DOI:** 10.1101/569772

**Authors:** Ana Cláudia Viana Wanzeler, Italo Antunes França Barbosa, Bruna Duarte, Eduardo Buzolin Barbosa, Daniel Almeida Borges, Monica Alves

**Affiliations:** Department of Ophthalmology, Faculty of Medical Sciences, University of Campinas - UNICAMP, Campinas, SP, Brazil.; Pontific Catholic University of Campinas – PUCCAMP, Campinas, SP, Brazil.

## Abstract

**Purpose:** To analyze how ocular surface parameters correlate to pterygium and investigate the possible impact on tear film and meibomian glands.

**Methods:** we investigated objective parameters of the ocular surface such as conjunctival hyperemia, tear film stability and volume, meibomian gland dysfunction, dry eye disease, corneal topography comparing healthy individuals and correlating with the pterygium clinical presentation.

**Results:** A total of 83 patients were included. Corneal astigmatism induction was 2.65 ± 2.52 D (0.4-11.8). The impact of pterygium on the ocular surface parameters compared to matched controls was seen in: conjunctival hyperemia (control 1.55±0.39/pterygium 2.14±0.69; p=0.0001), tear meniscus height (control 0.24±0.05 mm/pterygium 0.36±0.14mm; p 0.0002), meiboscore lower eyelid (control 0.29±0.64/pterygium 1.38±0.95; p 0.0001) and meiboscore upper eyelid (control 0.53±0.62/pterygium 0.98±0.75; p=0.0083). We found a high number of pterygium patients (88%) presented meibomian gland alterations. Interestingly, meibomian gland loss was coincident to the localization of the pterygium in 54% of the upper and 77% lower lids.

**Conclusion:** Pterygium greatly impacts on ocular surface by inducing direct alterations in the pattern of meibomian glands besides corneal irregularities, conjunctival hyperemia and lacrimal film alterations, inducing significant symptoms and potential signs of dysfunction.

## Introduction

Pterygium is a degenerative fibrovascular disease of the ocular surface that can cause symptoms of discomfort, corneal irregularities, aesthetics issues thus compromising visual acuity and patients’ quality of life. [1–3] The prevalence of pterygium varies worldwide. Global prevalence was estimated in 10.2% to 12%, reaching higher numbers in tropical regions. Several risk factors have been associated with pterygia, such as geographical latitude, residence in rural areas, old age, race, sex, sun exposure, chronic irritation and inflammation. [4,5]

Some studies have pointed tear film and ocular surface changes related to pterygium, but consistent correlations remain unknown. [6–9] Although numerous theories have been implicated in pterygium pathogenesis, including ultraviolet radiation exposure, viral infection, oxidative stress, genetic issues, inflammatory mediators, extracellular matrix modulators, it remains controversial as well as its impact on ocular surface homeostasis and function.[10] And a better understanding of the pathophysiological mechanisms associated with pterygium, the morphological alterations on the ocular surface and functional impact may contribute to specific approaches and more effective therapeutic proposals for this common ocular condition.

This study aimed to evaluate how ocular surface parameters correlate with pterygium clinical presentation and its impact on ocular surface structures and homeostasis.

## Materials and Methods

The present study had a transversal, observational and non-interventional design. It was performed after approval from the local research ethics committee and was conducted in accordance with the tenets of the Declaration of Helsinki and current legislation on clinical research. Written informed consent was obtained from all subjects after the explanation of the procedures and study requirements.

All propaedeutic methods were performed in accordance with specific guidelines and regulations. Data was collected during the ophthalmologic exams and in the inclusion of participants older than 18 years of age diagnosed with pterygium at the Cornea and External Disease Ambulatory, Department of Ophthalmology, University of Campinas.

Pterygium patients (n=52) and healthy volunteers (n=31) were included. We recorded personal and family history of pterygium, ocular and systemic comorbidities, ocular or systemic medications, visual acuity as well as a full ophthalmic exam. Ancilliary ocular surface evaluation consisted of: corneal topography, meibography, meniscometry, non-invasive tear film break-up time measurement, conjunctival hyperemia quantification using the Oculus Keratograph 5M (OCULUS Optikgerate GmbH, Wetzlar, Germany). All procedures were performed by the same examiner as detailed described below:

1. Tear film stability: evaluated the Non-invasive Tear Film Break-up Time (NITBUT) by Keratograph 5M through the evaluation of the point by point Placido concentric circles image during continuous eye-opening interval.
2. Tear meniscus height measured in millimeters in images taken by Keratograph 5M equipment.
3. Meibomian Gland Function: non-contact infrared meibography was performed in the lower and upper lid using Keratograph 5M. Gland dropout was assessed using meiboscan infrared device according to the instructions. Meiboscore was used for assessment of the meibography in the evaluation of the infrared captured images of the meibomian glands. The classification scale, adapted from Arita et al., used the following degrees for each eyelid: 0 (no loss of meibomian glands); 1 (loss of the meibomian gland involving less than one-third of the total meibomian gland area); 2 (loss between one third and two thirds of the total area of the meibomian gland); and 3 (loss more than two-thirds of the total meibomian gland area). [11]
4. Assessment of pterygium: pterygium pictures were used to classification in degrees: grade 1 to 4 according to fibrovascular tissue extension towards the cornea (grade 1 when the lesion reaches the limbus, grade 2 when it covers the cornea at about 2 mm, grade 3 when it reaches the pupil margin and grade 4 when it exceeds the pupil). Indeed, biomicroscopic aspect was noted as involutive atrophic or fleshy (involutive allows the visualization of structures immediately below and fleshy when fibrovascular tissue prevents proper visualization of underneath structures). [12] Hence, corneal topography images were taken for astigmatism measurements.

Exploratory data analysis was performed through summary measures (mean, standard deviation, minimum, median, maximum, frequency and percentage). Comparison between groups was performed using the Wilcoxon test. The correlation between numerical variables was assessed using the Spearman coefficient. The level of significance was 5%. The analyses were performed using the computer program The SAS System for Windows (Statistical Analysis System), version 9.4. (SAS Institute Inc, Cary, NC, USA).

## Results

A total of 83 patients were included in this study (52 pterygium patients and 31 healthy volunteers). Mean age of 53.69 ± 11.29 (26 – 75) years old in pterygium groups and 57.32 ± 7.30 (39–72) in healthy participants (p=0.6084).

Pterygia classification regarding tissue progression from limbus to the visual axis was: 1.9% as grade 1; 59.5% grade 2; 32.7% grade 3; and 5.8% as grade 4 (tissue over visual axis). In addition, 15.4% were atrophic and 84.6% had a fleshy/active clinical appearance. Corneal astigmatism induction was 2.65 ± 2.52 D (0.4-11.8). Table 1 shows ocular surface parameters in pterygium patients and controls and Table 2 shows data distribution according to the pterygium grades (a) and appearance (b).

**Table 1.** Comparisons of ocular Surface parameters in pterygium and healthy participants.

|  | Control |  | Pterygium |  | P value |
| --- | --- | --- | --- | --- | --- |
| | Mean ± SD | 95% CI | Mean ± SD | 95% CI | |
| Age (years) | 57.32±7.30 | 5.76 – 14.43 | 53.69±11.29 | 50.55 – 56.84 | 0.6084 |
| Visual Acuity | 0.85±0.21 | 0.77 – 0.94 | 0.63±0.31 | 0.54 – 0.71 | **0.0001\*** |
| Tear meniscus height | 0.24±0.05 | 0.22 – 0.26 | 0.36±0.14 | 0.32 – 0.40 | **0.0001\*** |
| NITBUT first Breakup | 10.6±6.51 | 8.22 – 13.01 | 10.64±5.29 | 9.16 – 12.11 | 0.8728 |
| NITBUT average Breakup | 14.28±6.06 | 11.88 – 16.68 | 13.55±5.55 | 12.01 – 15.10 | 0.6605 |
| Conjunctival hyperemia | 1.55±0.39 | 0.02-0.34 | 2.14±0.69 | 1.95-2.36 | **0.0001\*** |
| Meiboscore lower | 0.29±0.64 | 0.05 – 0.52 | 1.38±0.95 | 0.77 – 1.19 | **0.0001\*** |
| Meiboscore upper | 0.53±0.62 | 0.29 – 0.76 | 0.98±0.75 | 0.77 – 1.19 | **0.0083\*** |

**Table 2a.** Data distribution according to pterygium extension over the limbus (grades 0 – 4).

|  | Grades 1 – 2 |  | Grades 3 – 4 |  | P<br>value |
| --- | --- | --- | --- | --- | --- |
|  | Mean ± SD | 95% CI | Mean ± SD | 95% CI |  |
| Age (years) | 51.5 ± 11.3 | 47.3 – 55.6 | 57.1 ± 10.9 | 52 – 62.2 | 0.09 |
| K1 | 43.2 ± 2 | 42.5 – 44 | 42.8 ± 3.8 | 40.7 – 45 | 0.92 |
| K2 | 44.9 ± 1.9 | 44.1 – 45.6 | 47.5 ± 3.2 | 45.7 – 49.3 | **0.001\*** |
| Corneal astigmatism<br>(pterygium) | 1.6 ± 1.1 | 1.1 – 2 | 4.6 ± 3.3 | 2.7 – 6.4 | **0.006\*** |
| NITBUT | 9.8 ± 4.8 | 8.1 – 11.6 | 9.9 ± 5.1 | 7.6 – 12.3 | 0.95 |
| Conjunctival<br>hyperemia | 2.7 ± 0.6 | 2.4 – 2.9 | 2.7 ± 0.6 | 2.4 – 3 | 0.96 |
| Tear meniscus height | 0.3 ± 0.1 | 0.3 – 0.4 | 0.3 ± 0.1 | 0.2 – 0.4 | 0.27 |
| Red eye (0-10) | 7.7 ± 3 | 6.3 – 9.1 | 8 ± 2.1 | 6.7 – 9.4 | 0.96 |
| Irritation (0-10) | 7.1 ± 1.7 | 6.3 – 7.9 | 6.5 ± 3.4 | 4.3 – 8.6 | 0.92 |
| Tearing (0-10) | 5.8 ± 3.6 | 4.1 – 7.4 | 6 ± 3.9 | 3.4 – 8.5 | 0.76 |
| Blurred vision (0-10) | 6.7 ± 3.5 | 5.1 – 8.3 | 7.2 ± 2.9 | 5.4 – 9 | 0.95 |
| Aesthetics (0-10) | 7.9 ± 3 | 6.5 – 9.3 | 7.5 ± 3.2 | 5.4 – 9.5 | 0.66 |

**Table 2b.** Data distribution according to the pterygium clinical presentation (involutive and fleshy).

|  | Atrophic |  | Fleshy |  | P<br>value |
| --- | --- | --- | --- | --- | --- |
|  | Mean ± SD | 95% CI | Mean ± SD | 95% CI |  |
| Age (years) | 58.5 ± 13.3 | 47.3 – 69.6 | 52.8 ± 10.8 | 49.5 –<br>56.1 | 0.26 |
| K1 | 42.7 ± 3.3 | 40 – 45.5 | 43.2 ± 2.5 | 42.3 – 44 | 0.40 |
| K2 | 46.2 ± 5.2 | 41.8 – 50.6 | 45.7 ± 1.8 | 45 – 46.2 | 0.40 |
| Corneal astigmatism<br>(pterygium) | 3.5 ± 2.7 | 1.1 – 5.8 | 2.4 ± 2.4 | 1.6 – 3.2 | 0.18 |
| NITBUT | 9.5 ± 7.1 | 3.5 – 15 | 10 ± 4.5 | 8.7 – 11.5 | 0.36 |
| Conjunctival<br>hyperemia | 2.3 ± 0.1 | 2.5 – 2.9 | 2.7 ± 0.7 | 1.8 – 2.9 | **0.005\*** |
| Tear meniscus height | 0.4 ± 0.2 | 0.2 – 0.6 | 0.3 ± 0.1 | 0.31 –<br>0.39 | 0.25 |
| Red eye (0-10) | 9 ± 1 | 7.7 – 10 | 7.7 ± 2.8 | 6.6 – 8.8 | 0.82 |
| Irritation (0-10) | 7 ± 2 | 4.5 – 9.4 | 6.8 ± 2.5 | 5.9 – 7.8 | 0.51 |
| Tearing (0-10) | 6.4 ± 2.6 | 3 – 9.6 | 5.8 ± 3.8 | 4.4 – 7.3 | 0.92 |
| Blurred vision (0-10) | 8.8 ± 1 | 7.4 – 10 | 6.7 ± 3.4 | 5.4 – 7 | 0.25 |
| Aesthetics (0-10) | 7 ± 2.9 | 3.3 – 10.6 | 8 ± 3 | 6.8 – 9 | 0.34 |

Compared to control participants, pterygium patients presented significant alterations regarding hyperemia (control 1.55±0.39 - 95% CI 0.02-0.34; pterygium 2.14±0.69 - 95% CI 1.95-2.36; p 0.0001), tear meniscus height (control 0.24±0.05 mm - 95% CI 0.22-0.26; pterygium 0.36±0.14mm - 95% CI 0.32-0,40; p 0.0002) and meiboscore lower eyelid (control 0.29±0.64 - 95% CI 0.05-0.52; pterygium 1.38±0.95 - 95% CI 0.77-1.19; p 0.0001) and meiboscore upper eyelid (control 0.53±0.62– 95% IC 0.29-0.76; pterygium 0.98±0.75 - 95% IC 0.77-1.19; p 0.0083).

We found that 88% of patients presented abnormalities on meibomian glands. Interestingly, in 54% of the upper eyelids and 77% of the lower eyelids, the meibomian gland loss appeared nasally in the same localization of the pterygium. Figure 1 shows the distribution of the meibomian gland involvement in the upper and lower eyelid and Figure 2 exemplifies the meibography alterations in pterygium patients.

**Figure 1:**
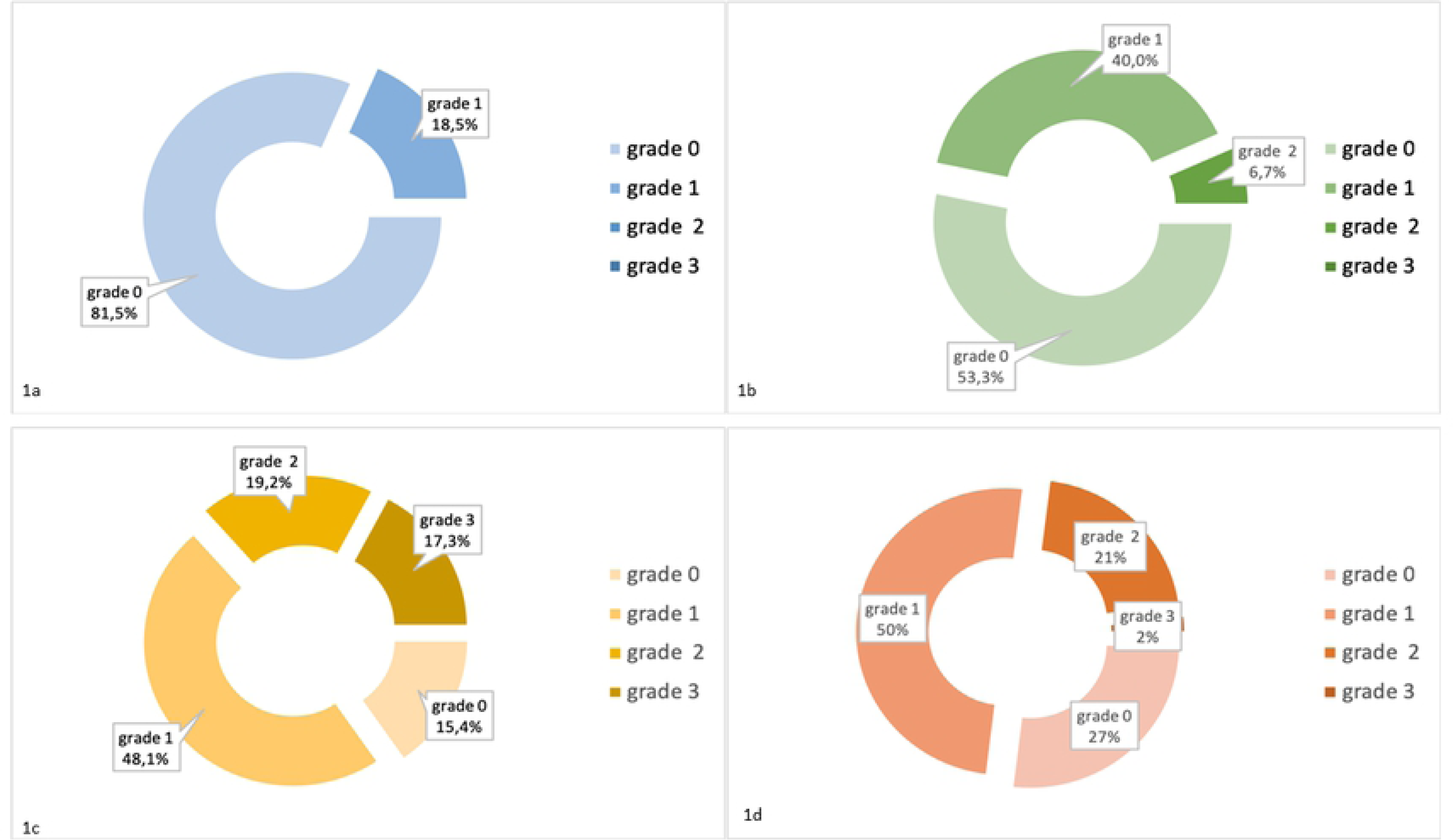
Meibomian gland evaluation in control individuals (1a: lower eyelid; 1b: upper eyelid) and pterygium patients (1c: lower eyelid; 1d: upper eyelid). Meiboscore grade (0-3)

**Figure 2:**
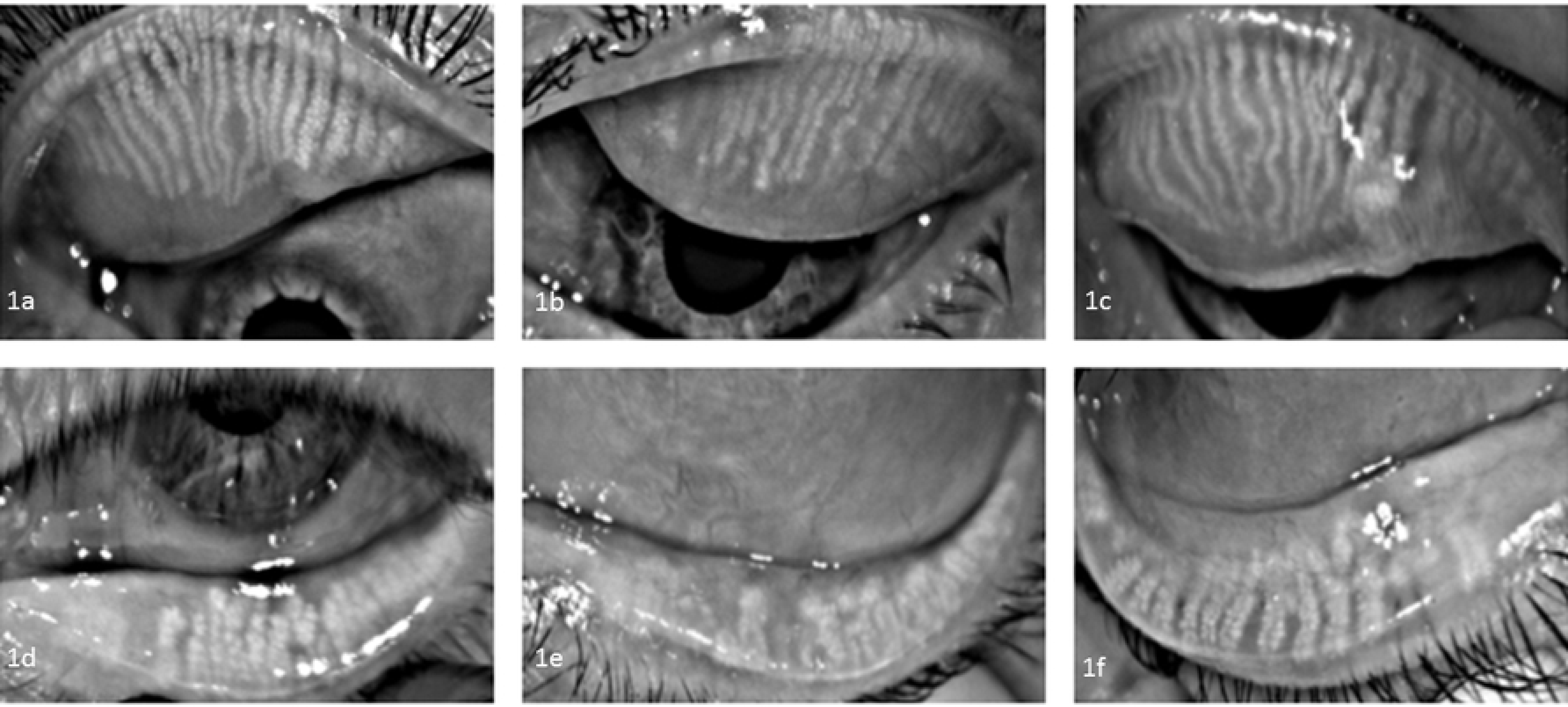
Meibomian gland dysfunction in pterygium patients: nasal pterygium and upper eyelid dropout (1a and 1b right eyes and 1c left eye) and nasal pterygium and lower eyelid dropout (1d and 1e right eyes and lf left eye).

Regarding the subjective symptoms, the patients’ complaints were evaluated as parameters such as tearing, ocular discomfort, aesthetics and blurred vision. Such symptoms are closely related to the tear film and ocular surface abnormalities described.

## Discussion

The present study shows that pterygium has a great impact on the parameters and structures of the ocular surface. It can induce corneal astigmatism, conjunctival hyperemia, tear film abnormalities and significant structural alterations in the meibomian glands.

Ocular hyperemia can be considered as a clinical sign of inflammation that may suggest severity and progression of a specific disease. [13] High rates of hyperemia were observed in the eyes with pterygium, which may be explained by the number of fleshy pterygia present in this study and by the richer vascularization of the pterygium itself, even in the atrophic ones. The advantage of using the image analysis method is that one can eliminate individual variability and the bias of subjective classification. [14]

Although pterygium symptoms resemble dry eye and others ocular surface diseases symptoms, such as dryness and irritation, no decrease on non-invasive tear break-up time (NITBUT) was observed in this cohort. A study carried out in 2014 had already shown that the size of the pterygium does not correlate with the tear break-up time and the results of the Schirmer’s test. [15] Another study comparing Schirmer’s test results and tear break-up time before and after pterygium surgery showed that, even with the removal of the pterygium, there were no changes in those tests results one month after surgery. [16] On the other hand, Ozsutu et al. found lower values of tear film test and Schirmer I test in eyes with pterygium when compared to healthy eyes, which can be explained by the significantly higher tear osmolarity levels found in the study.[6] In our study pterygium patients had statistically higher tear meniscus height and no differences in NITBUT, which may be pointed as a picture of the ocular surface compensatory mechanisms. Higher measurements of tear meniscus may be related to chronic ocular inflammation and friction and abnormal distribution of tear film leading to surface disturbances in tear flow dynamics and reflex tearing, as described in previous studies.[16] Although, normal tear function and no alteration on tear meniscus height has been already described in the pterygium patients [14,17]

Pterygium can induce corneal aberrations that compromise patients’ visual acuity. Studies indicate the length of the pterygium and vascularization as predictive factors for increased induction of astigmatism.[17–19] On the other hand, pterygium excision leads to a decrease in acquired astigmatism to acceptable levels, as shown by studies that evaluated the impact of surgery on corneal astigmatism reduction.[20] Regarding the surgical procedure, there was no significant effect on the degree of astigmatism were found comparing different surgical techniques.[21]

Meibomian gland dysfunction was found in a significant number of pterygium patients. Interestingly, areas of meibomian gland loss coincidently correlated to the nasal topography localization of the pterygium, both in the upper and lower eyelids. Another study reported morphological changes in such pterygium-associated glands when compared to healthy eyes, demonstrating that patients with pterygium had a significantly higher degree of meibomian gland loss when compared to normal patients, but to our knowledge, no association with pterygium localization was described before. [22] We hypothesized that those changes might be related to local inflammatory conditions and the release of inflammatory cytokines that can spread to the anterior and posterior margin of the eyelid, resulting in meibomian gland alterations, as seen in other ocular surface disorders.[23] Indeed, an effect of the direct friction caused by the pterygium in the tarsal conjunctiva may play a contributory role. However, this finding demands further exploration. Of note, meibomian gland proper production and delivery is crucial to tear film stability and evaporative dry eye is considered the most common subtype of disease affection a great number of individuals worldwide. Thus, disruption of meibomian gland function negatively impacts both the quality and quantity of meibum and in turn affects ocular surface health. Increased tear evaporation, tear film instability and consequent hyperosmolarity, inflammation and ocular surface damage lead to ocular discomfort and visual disruption. The findings related to pterygium and meibomian gland described in this study, may indicate a need of closer attention regarding to quantification of related symptoms, search of clinical signs and overall preventive measurements to guarantee meibomian gland functional support, such as eyelid hygiene, mechanical expression and other procedures.

Some limitations of this study must be pointed out. Our sample consisted of patients that consecutively presented for consultation. Although with distinct grades according to the proposed classification system, the included participants had primary, nasal pterygia. Recurrent and temporally located pterygia may carry different features, not evaluated in this study. The use of noninvasive technology for ocular surface study has proved to be of great value, but a broad investigation of tear and tissue inflammatory mediators may enhance the understanding of pterygium mechanisms and the ocular surface changes described herein.

This study demonstrated a detailed evaluation of the clinical parameters of the ocular surface in the pterygium population and quantified the symptoms. Therefore, our results not only allowed for a contribution to the understanding of the disease but also created new perspectives for future studies.

## Conclusion

The present study shows that pterygium impacts on ocular surface parameters, especially by inducing direct alterations in the pattern of meibomian glands and tear film.

## References

1. Saw SM, Tan D. Pterygium: prevalence, demography and risk factors. Ophthalmic epidemiology. 1999 Sep;6(3):219–28.

2. Chui J, Girolamo N Di, Wakefield D, Coroneo MT. The Pathogenesis of Pterygium: Current Concepts and Their Therapeutic Implications. The Ocular Surface [Internet]. 2008 Jan 1 [cited 2018 Mar 18];6(1):24–43. Available from: https://www.sciencedirect.com/science/article/pii/S1542012412701039

3. Chui J, Coroneo MT, Tat LT, Crouch R, Wakefield D, Di Girolamo N. Ophthalmic pterygium: A stem cell disorder with premalignant features. American Journal of Pathology [Internet]. 2011 Feb;178(2):817–27. Available from: http://dx.doi.org/10.1016/j.ajpath.2010.10.037

4. Liu L, Wu J, Geng J, Yuan Z, Huang D. Geographical prevalence and risk factors for pterygium: a systematic review and meta-analysis. BMJ Open [Internet]. 2013/11/19. 2013;3(11):e003787. Available from: https://www.ncbi.nlm.nih.gov/pubmed/24253031

5. Rezvan F, Khabazkhoob M, Hooshmand E, Yekta A, Saatchi M, Hashemi H. Prevalence and risk factors of pterygium: a systematic review and meta-analysis. Survey of ophthalmology. 2018 Sep;63(5):719–35.

6. Ozsutcu M, Arslan B, Erdur SK, Gulkilik G, Kocabora SM, Muftuoglu O. Tear osmolarity and tear film parameters in patients with unilateral pterygium. Cornea. 2014 Nov;33(11):1174–8.

7. Ergin A, Bozdoğan Ö. Study on Tear Function Abnormality in Pterygium. Ophthalmologica [Internet]. 2001;215(3):204–8. Available from: https://www.karger.com/DOI/10.1159/000050859

8. Chan CML, Liu YP, Tan DTH. Ocular surface changes in pterygium. Cornea. 2002 Jan;21(1):38–42.

9. Wu H, Lin Z, Yang F, Fang X, Dong N, Luo S, et al. Meibomian Gland Dysfunction Correlates to the Tear Film Instability and Ocular Discomfort in Patients with Pterygium. Scientific Reports [Internet]. 2017 Mar 24;7:45115. Available from: https://doi.org/10.1038/srep45115

10. Bradley JC, Yang W, Bradley RH, Reid TW, Schwab IR. The science of pterygia. The British journal of ophthalmology. 2010 Jul;94(7):815–20.

11. Arita R, Suehiro J, Haraguchi T, Shirakawa R, Tokoro H, Amano S. Objective image analysis of the meibomian gland area. The British Journal of Ophthalmology [Internet]. 2014 Jun 27;98(6):746–55. Available from: http://www.ncbi.nlm.nih.gov/pmc/articles/PMC4033206/

12. Schellini SA, dos Reis Veloso CE, Lopes W, Padovani CR, Padovani CRP. Characteristics of patients with pterygium in the Botucatu region. Arquivos brasileiros de oftalmologia. 2005;68(3):291–4.

13. Murphy PJ, Lau JSC, Sim MML, Woods RL. How red is a white eye? Clinical grading of normal conjunctival hyperaemia. Eye (London, England). 2007 May;21(5):633–8.

14. Hilmi MR, Che Azemin MZ, Mohd Kamal K, Mohd Tamrin MI, Abdul Gaffur N, Tengku Sembok TM. Prediction of Changes in Visual Acuity and Contrast Sensitivity Function by Tissue Redness after Pterygium Surgery. Current eye research. 2017 Jun;42(6):852–6.

15. Kampitak K, Leelawongtawun W. Precorneal tear film in pterygium eye. Journal of the Medical Association of Thailand = Chotmaihet thangphaet. 2014 May;97(5):536–9.

16. Kampitak K, Tansiricharernkul W, Leelawongtawun W. A comparison of precorneal tear film pre and post pterygium surgery. Journal of the Medical Association of Thailand = Chotmaihet thangphaet. 2015 Mar;98 Suppl 2:S53-5.

17. Han SB, Jeon HS, Kim M, Lee S-J, Yang HK, Hwang J-M, et al. Quantification of Astigmatism Induced by Pterygium Using Automated Image Analysis. Cornea. 2016 Mar;35(3):370–6.

18. Avisar R, Loya N, Yassur Y, Weinberger D. Pterygium-induced corneal astigmatism. The Israel Medical Association journal : IMAJ. 2000 Jan;2(1):14–5.

19. Oner FH, Kaderli B, Durak I, Cingil G. Analysis of the Pterygium Size Inducing Marked Refractive Astigmatism. European Journal of Ophthalmology [Internet]. 2000 Jul 1;10(3):212–4. Available from: https://doi.org/10.1177/112067210001000304

20. Khan FA, Khan Niazi SP, Khan DA. The impact of pterygium excision on corneal astigmatism. Journal of the College of Physicians and Surgeons--Pakistan : JCPSP. 2014 Jun;24(6):404–7.

21. Altan-Yaycioglu R, Kucukerdonmez C, Karalezli A, Corak F, Akova YA. Astigmatic changes following pterygium removal: comparison of 5 different methods. Indian journal of ophthalmology. 2013 Mar;61(3):104–8.

22. Ye F, Zhou F, Xia Y, Zhu X, Wu Y, Huang Z. Evaluation of meibomian gland and tear film changes in patients with pterygium. Indian journal of ophthalmology. 2017 Mar;65(3):233–7.

23. Ibrahim OMA, Matsumoto Y, Dogru M, Adan ES, Wakamatsu TH, Shimazaki J, et al. In vivo confocal microscopy evaluation of meibomian gland dysfunction in atopic-keratoconjunctivitis patients. Ophthalmology. 2012 Oct;119(10):1961–8.

